# Cancer Stem Cell Enrichment and Metabolic Substrate Adaptability are Driven by Hydrogen Sulfide Suppression in Glioblastoma

**DOI:** 10.1101/2020.03.08.982116

**Authors:** Daniel J. Silver, Gustavo A. Roversi, Nazmin Bithi, Chase K. A. Neumann, Katie M. Troike, Grace K. Ahuja, Ofer Reizes, J. Mark Brown, Christopher Hine, Justin D. Lathia

**Affiliations:** Cancer Impact Area and Lerner Research Institute, Cleveland Clinic Foundation, Cleveland, Ohio, USA; Department of Cardiovascular and Metabolic Sciences, Lerner Research Institute, Cleveland Clinic Foundation, Cleveland, Ohio, USA; Cleveland Clinic Lerner College of Medicine, Cleveland Clinic Foundation, Cleveland, Ohio, USA; Case Comprehensive Cancer Center, Case Western Reserve University, Cleveland, Ohio, USA; Rose Ella Burkhardt Brain Tumor and Neuro-Oncology Center, Cleveland Clinic, Cleveland, Ohio, USA

**Keywords:** Glioblastoma, Cancer Stem Cells, Metabolism, Diet, Hydrogen Sulfide

## Abstract

Glioblastoma (GBM) remains among the deadliest of human malignancies. The emergence of the cancer stem cell (CSC) phenotype represents a major challenge to disease management and durable treatment response. The extrinsic, environmental, and lifestyle factors that result in CSC enrichment are not well understood. The CSC state endows cells with a fluid metabolic profile, enabling the utilization of multiple nutrient sources. Therefore, to test the impact of diet on CSC enrichment, we evaluated disease progression in tumor-bearing mice fed an obesity-inducing high-fat diet (HFD) *versus* an energy-balanced, low-fat control diet. HFD consumption resulted in hyper-aggressive disease that was accompanied by CSC enrichment and shortened survival. HFD consumption also drove intracerebral accumulation of saturated fats, which in turn inhibited the production and signaling of the gasotransmitter hydrogen sulfide (H_2_S). H_2_S is an endogenously produced bio-active metabolite derived from sulfur amino acid catabolism. It functions principally through protein S-sulfhydration and regulates a variety of programs including mitochondrial bioenergetics and cellular metabolism. Inhibition of H_2_S synthesis resulted in increased proliferation and chemotherapy resistance, whereas treatment with H_2_S donors led to cytotoxicity and death of cultured GBM cells. Compared to non-cancerous controls, patient GBM specimens were reduced in overall protein S-sulfhydration, which was primarily lost from proteins regulating cellular metabolism. These findings support the hypothesis that diet-regulated H_2_S signaling serves to suppress GBM by restricting metabolic adaptability, while its loss triggers CSC enrichment and disease acceleration. Interventions augmenting H_2_S bioavailability concurrent with GBM standard of care may improve outcomes for GBM patients.

**One Sentence Summary:** Consumption of a high-fat diet (HFD) accelerates glioblastoma (GBM) by inhibiting the production and signaling of the tumor-suppressive metabolite hydrogen sulfide (H_2_S).

## Introduction

Patients with glioblastoma (GBM) experience one of the most aggressive disease trajectories of all cancers*(1*, *2)*. Effective disease management is lacking due to the intrinsic heterogeneity and complexity of the disease. Heterogeneity at the cellular level includes multiple interacting populations of cancer stem cells (CSCs)*(3–5)*, each capable of varying degrees of tissue invasion*(6)*, proliferation*(7*, *8)*, and treatment resistance*(9)*. New evidence indicates that metabolic heterogeneity confers an additional level of complexity and adaptability to evolving GBM tumors*(10)*. A growing number of enzymes and metabolic pathways have been highlighted that contribute to the maintenance and selection of CSC populations*(11*–*13)*. Collectively, these studies suggest that metabolic adaptability enables CSCs to out-compete less-plastic tumor cells with clear metabolic dependencies. Specifically, while non-stem tumor cells may be limited to the glycolytic metabolism first described by Otto Warburg, the CSC phenotype enables cells to shift among various substrates, employing glycolysis, fatty acid oxidation, and amino and nucleic acid metabolism depending on nutrient availability*(14)*. As the nutrient landscape is profoundly altered through diet, we focused on understanding how an obesity-generating, high-fat diet (HFD) serves as a regulator of GBM progression and as a selective force for GBM CSC expansion.

HFD consumption alters numerous physiological systems, including the lipid composition within the brain*(15)* as well as the composition of the gut microbiome*(16*, *17)* and its associated set of metabolites. HFD also modifies the composition and function of the immune system*(18)*. One system that is profoundly affected by HFD consumption and has been largely unexplored in the GBM literature is the synthesis of the gasotransmitter hydrogen sulfide (H_2_S), a byproduct of sulfur amino acid metabolism. H_2_S is an endogenously produced, bio-active metabolite*(19)*. Three enzymes, cystathionine beta-synthase (CBS), cystathionine gamma-lyase (CGL), and mercaptopyruvate sulfurtransferase (MPST), are responsible for enzymatic H_2_S production and are differentially expressed and active in a tissue-specific manner. Through a largely unknown mechanism, HFD consumption results in attenuation of these H_2_S producing enzymes and therefore potently inhibits H_2_S production*(20)*. Functionally, upon generation, H_2_S is quickly transferred to available cysteine residues in the form of a protein post-translational modification referred to as S-sulfhydration*(21)*. S-sulfhydration alters protein form and associated function; however, unlike nitrosylation or phosphorylation, the functional changes that result from S-sulfhydration are largely unknown*(22)*. There is limited information on H_2_S and cancer*(23*–*25)*. Evidence suggests that H_2_S serves as both a promoter and inhibitor of tumorigenesis in a tissue-specific manner. For GBM, the limited information available suggests that H_2_S synthesis inhibits proliferation of cultured GBM tumor cells*(26)*. Biochemical analysis indicates that the enzymatic activity of both CGL and MPST decreases with increasing astrocytoma grade and that these enzymes are essentially non-functioning in the context of GBM*(27)*. To date, the loss of S-sulfhydration has gone entirely unexplored in the context of GBM.

HFD has been the subject of two types of research in the GBM field. First, metabolic dependency studies involve the use of ketogenic HFDs or metabolism pathway inhibitors to slow or stop disease progression by depriving tumors of critical energy sources*(28*–*30)*. Second, epidemiological studies question whether obesity, brought about by the consumption of obesity-generating diets, serves as an initiator of GBM development*(31*, *32)*. The possibility that a long-term pattern of HFD consumption would exacerbate disease, changing the histological presentation and trajectory of GBM, has not yet been addressed. Thus, we compared GBM tumors developed in the brains of HFD-fed mice to those developed within the brains of animals fed a control diet. This led to a series of connected observations linking HFD consumption to alterations in the nutrient landscape of the brain, the CSC compartment of the tumor microenvironment, and the production and function of intracerebral H_2_S.

## Results

### High-Fat Diet Consumption Drives CSC Enrichment and Accelerates Glioblastoma Progression

To test whether HFD consumption modulates the growth and initiation of GBM, we employed both syngeneic mouse models (GL261, CT2A, KR158) and a human patient-derived GBM model (*h*GBM 23) in a series of *in vivo* experiments performed according to the schematic presented in **Figure 1A**. Experiments were initiated using animals of equivalent body mass and fat composition. Animals fed the HFD gained body mass (**Supplemental Figure 1A**) as a product of fat accumulation (**Supplemental Figure 1B**) over time throughout the duration of the experiment. Compared to control diet-fed mice, HFD-fed tumor-bearing animals experienced a significant reduction of overall survival (**Figure 1B – D; Supplementary Figure 1C**). Importantly, in the absence of GBM, HFD consumption does not limit survival. Under specific experimental conditions, HFD has even been attributed to increased longevity*(33)* and protection against midlife mortality*(34)* in rodent models of aging. For each of the three syngeneic GBM models, these experiments were repeated across multiple cohorts in a limiting-dilution format using progressively fewer tumor cells at the time of intracerebral injection. Regardless of the initial cell dosage, a greater number of animals in the HFD group succumbed to disease in the time allotted to the experiment, indicating that HFD consumption induced a higher tumor initiation frequency compared to consumption of the control diet. Specifically, 2-3-fold fewer tumor cells were required to initiate tumors that drove animals to the experimental endpoint under conditions of HFD consumption (**Figure 1E – G**). Because we interpreted this enhanced tumor aggression as indicating CSC enrichment, we assessed end-stage tumors using standard immunofluorescence techniques for the CSC-associated transcription factor SOX2. In accordance with the increased CSC frequency suggested by the *in vivo* limiting-dilution analysis, histological examination revealed a marked increase in the percentage of SOX2^+^ tumor cells within the brains of HFD-fed mice compared to mice fed the control diet (**Figure 1H – J**). Thus, consumption of a high-fat, obesogenic diet resulted in a hyper-aggressive GBM phenotype and associated enrichment of CSCs within the tumors of HFD-fed mice.

**Figure 1.**
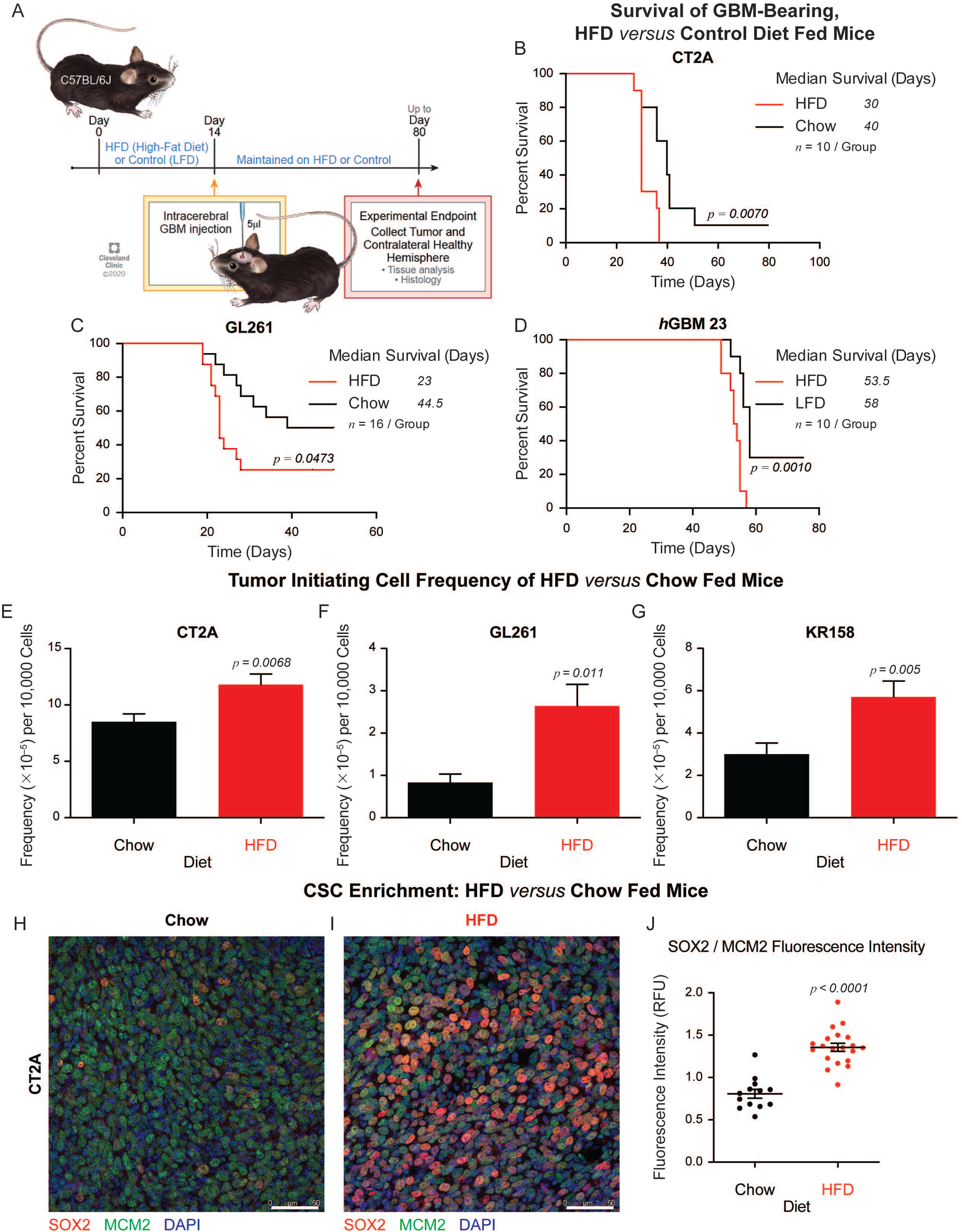
High-Fat Diet Consumption Drives CSC Enrichment and Accelerates Glioblastoma Progression. (**A**) *In vivo* experimental design employed to test whether HFD consumption modifies GBM progression. (**B – D**) For the syngeneic GBM models CT2A and GL261, as well as the patient-derived GBM model *h*GBM 23, Kaplan-Meier survival analysis confirmed significant truncation of overall survival under conditions of HFD consumption compared to consumption of control diets. *p* values determined by log-rank survival analysis and experimental group (*n*) size noted above. (**E – G**) *In vivo* limiting dilution analysis was performed for the three syngeneic GBM models CT2A, GL261, and KR158. For each model, tumors were initiated using 50,000, 20,000, 15,000, 10,000, and 5,000 cells/animal. *p* values were determined using the Walter and Eliza Hall ELDA portal*(52)* comparing the total number of endpoint animals in the HFD group *versus* the control diet group at the conclusion of each set of experiments. (**H, I**) Representative immunofluorescent micrographs of the CSC population observed in the GBM tumor microenvironment under HFD- *versus* chow-fed conditions. The CSC-associated transcription factor SOX2 was visualized in red; MCM2, visualized in green, identified the bulk tumor cell population; and nuclei were visualized in blue using DAPI. (**J**) SOX2 fluorescence intensity, normalized to the MCM2 fluorescence intensity, allowed us to measure CSC enrichment within the tumor microenvironment. *p* value determined by unpaired t-test.

### HFD Consumption Drives Stem Cell Phenotype Enrichment

We reasoned that HFD consumption may result in intracerebral lipid enrichment, which in turn may act directly (and/or indirectly) to increase proliferation and self-renewal within the tumor cell population. Therefore, matched tumor-bearing and non-tumor-bearing hemispheres were isolated from the brains of multiple HFD-fed and control diet-fed GBM-injected animals at their experimental endpoints. To determine which lipid species were altered, we interrogated these specimens using mass spectrometry-based non-targeted lipidomic analysis (**Figure 2A**, **Table 1**). When normalized to the set of chow-fed healthy specimens, we observed two modes of *in vivo* lipid alteration. First, we identified a variety of lipids enriched specifically within the tumors isolated from HFD-fed mice (**Figure 2B**). These lipid species were expressed at significantly greater levels within tumors of HFD-fed animals compared to the contralateral healthy tissues derived from the same mice as well as tissues isolated from control diet-fed mice. Second, we identified a separate set of lipids enriched generally within the brains of HFD-fed animals, regardless of the presence of tumor (**Figure 2C**). Expression of these species was significantly increased in the HFD-derived specimens compared to the control diet-fed specimens but was not different between the tumor-derived and contralateral specimens isolated from the HFD-fed mice. With the exception of two polyunsaturated ceramide species (HexCer 34:1;2 and HexCer 34:1;3), the lipids generated within the HFD-fed, tumor-bearing brain were saturated, mono-, or di-unsaturated lipid species, consistent with heavy consumption of a saturated fat-based diet. As ceramide lipids have a strong association with cellular stress*(35*, *36)*, which is commonly induced within the caustic growth zones*(37*, *38)* of GBM, we omitted these species from our analysis. Based on this non-targeted lipid assessment, we concluded that HFD consumption induced accumulation of saturated fats within the brain and tumor microenvironment of HFD-fed animals. This novel lipid accumulation, in turn, may have contributed to the hyper-aggressive disease that presented in these animals. Rather than narrow our study to the function of an individual saturated lipid, we hypothesized that this overall saturated lipid accumulation drove increased proliferation, CSC phenotype induction, and/or CSC selection and expansion.

**Figure 2.**
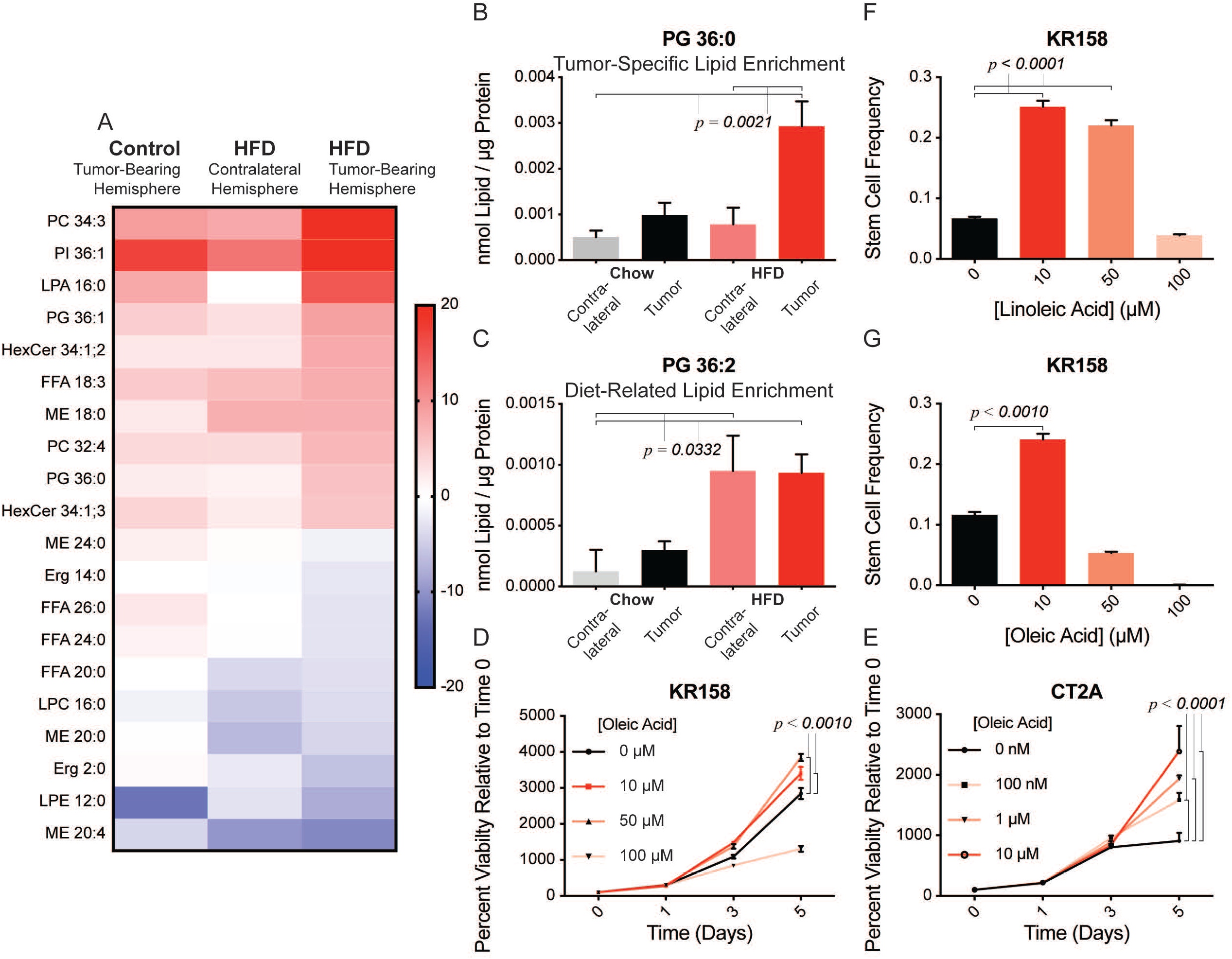
Consumption of HFD Drives Intracerebral Lipid Accumulation, Promoting Tumor Cell Viability and Self-Renewal. (**A**) Heatmap representing the top 10 most abundant and the bottom 10 least abundant lipids out of 216 total lipid species identified by untargeted lipidomic analysis. Four groups (*i*. HFD-fed, tumor-bearing hemisphere, *ii*. HFD-fed, contralateral hemisphere, *iii*. Chow-fed tumor-bearing hemisphere, and *iv*. Chow-fed, contralateral hemisphere) were compared; *n* = 5 specimens/group. Heatmap data were normalized to group *iv*, the Chow-fed, contralateral hemisphere group. Two modes of lipid enrichment were noted. (**B**) Four saturated lipid species, including phosphatidylglycerol (PG) 36:0, were identified specifically within the tumors of the HFD-fed animals. *p* value determined by one-way ANOVA. (**C**) Nine mono- or di-unsaturated lipid species, including phosphatidylglycerol (PG) 36:2, were identified within the HFD-fed brain and tumor. *p* value determined by one-way ANOVA. *In vitro* treatment of either (**D**) KR158 or (**E**) CT2A with the mono-unsaturated fatty acid oleic acid 18:1 increased tumor cell viability in a dose-dependent manner. *p* value determined by two-way ANOVA; 5 technical replicates per timepoint. (**F, G**) *In vitro* limiting dilution analysis conducted with KR158 indicated self-renewal enhancement resulting from exposure to excess oleic or linoleic acid. *p* values were determined using the Walter and Eliza Hall ELDA portal*(52)*.

To test whether excess saturated lipid might increase tumor cell proliferation and self-renewal, we supplemented the standard growth media employed for the syngeneic GBM models with the mono- or di-unsaturated fatty acids oleic or linoleic acid. We then compared cellular growth and self-renewal in the presence of exogenous fatty acid to growth under control conditions. GBM cells grown in excess lipid were induced into a state of hyper proliferation (**Figure 2D, E**) and exhibited increased self-renewal (**Figure 2F, G**). Thus, in accordance with our hypothesis, we concluded that excess saturated lipid was acting directly on tumor cells and contributing to the enhanced GBM progression and increased CSC frequency observed in the context of HFD consumption.

Importantly, while these findings indicated that saturated fats may work directly on tumor cells, they did not rule out the possibility that lipid accumulation within the HFD-fed brain may have established an environment that facilitated outgrowth of a stem-like population. Our lipidomic profiling revealed a host of species that accumulated in the brains of HFD-fed animals regardless of the presence of a tumor. This finding reinforced the idea that long-term HFD consumption might shift the overall nutrient landscape of the brain, thus serving as a selective pressure for the enrichment of stem-like cells with an enhanced ability to forage and metabolize a diverse set of energy substrates, including lipids *(39)*. To test whether HFD consumption established a stem cell-selective environment, we introduced a cohort of female, non-tumor-bearing C57BL/6J mice to *ad libitum* HFD, matched to a control cohort maintained on standard rodent chow. These differential diets were maintained for approximately 50 days, a time period roughly equal to the survival of HFD-fed GBM-transplanted mice. Using standard immunofluorescence techniques, we then carefully examined the subventricular zones (SVZs) of these differentially fed animals for the expansion of endogenous neural stem and progenitor cells (NSPCs). Staining for the stem cell-associated transcription factor SOX2 revealed amplification of the NSPC fraction within the SVZs of the HFD-fed cohort compared to control animals (**Supplemental Figure 2**). Based on these data, we concluded that HFD consumption established an environment within the brain in which proliferation and self-renewal of stem-like tumor and NSPCs was selectively enhanced. Therefore, we proceeded to evaluate molecular mechanisms that are holistically modulated by HFD.

### The Diet-Modifiable Metabolite Hydrogen Sulfide Serves as a GBM Tumor Suppressor

Long-term HFD consumption inhibits production of the gasotransmitter H_2_S, a byproduct of cysteine metabolism and a feature of the transsulfuration metabolic pathway (**Figure 3A**). While this diet-induced inhibition has been well documented in the livers of HFD-fed mice*(20)*, we observed an approximate 50% reduction in H_2_S synthesis within the brains of HFD-fed tumor-bearing mice (**Figure 3B**). We began our investigation into H_2_S and GBM by analyzing transcript expression data curated by The Cancer Genome Atlas to ascertain whether H_2_S synthesizing enzymes might have prognostic value for patients with GBM. Correlating the mRNA expression of the three H_2_S-synthesizing enzymes with overall survival indicated that patients presenting low expression of CBS and MPST experienced truncated survival compared to patients in which H_2_S production remained intact (**Figure 3C, D**). While the mRNA expression level of CGL was not prognostic (**Figure 3E**), recent work confirmed that the enzymatic function of both MPST and CGL was suppressed in the context of GBM*(27)*. These findings suggest a tumor-suppressive role for H_2_S insofar as the shutdown of H_2_S synthesis confers a growth advantage to GBM tumors*(26)*. To test the hypothesis that H_2_S serves as a GBM tumor suppressor, we assessed the proliferation of cultured GBM cells treated with the potent and selective CGL inhibitor propargylglycine (PAG). Treatment with PAG inhibited H_2_S production in each of the syngeneic GBM models (**Supplemental Figure 3A**). Further, inhibition of H_2_S synthesis induced hyperproliferation (**Figure 3F – H**) and protected against the cytotoxic effects of the standard-of-care chemotherapeutic temozolomide compared to vehicle controls (**Supplementary Figure 3B**). Because inhibition of H_2_S synthesis drove GBM cell proliferation, we reasoned that H_2_S replacement should suppress GBM tumor cell growth. We compared the *in vitro* IC_50_ values for sodium hydrosulfide (NaHS) and GYY4137, two H_2_S donors, using multiple syngeneic and patient-derived GBM models, as well as two liver cancer control cell lines (HepG2 and NCTC) (**Figure 3I – K**, **Supplementary Figure 3C**). We observed that cultured GBM cell (human or mouse) viability was suppressed to a far greater degree than that of either of the control liver cancer cell lines. These data support the conclusion that H_2_S is a diet-modifiable tumor suppressor of GBM. Additionally, these data suggest that HFD consumption sufficiently depleted this tumor suppressor, such that the HFD-fed mice experienced a hyper-aggressive version of the disease.

**Figure 3.**
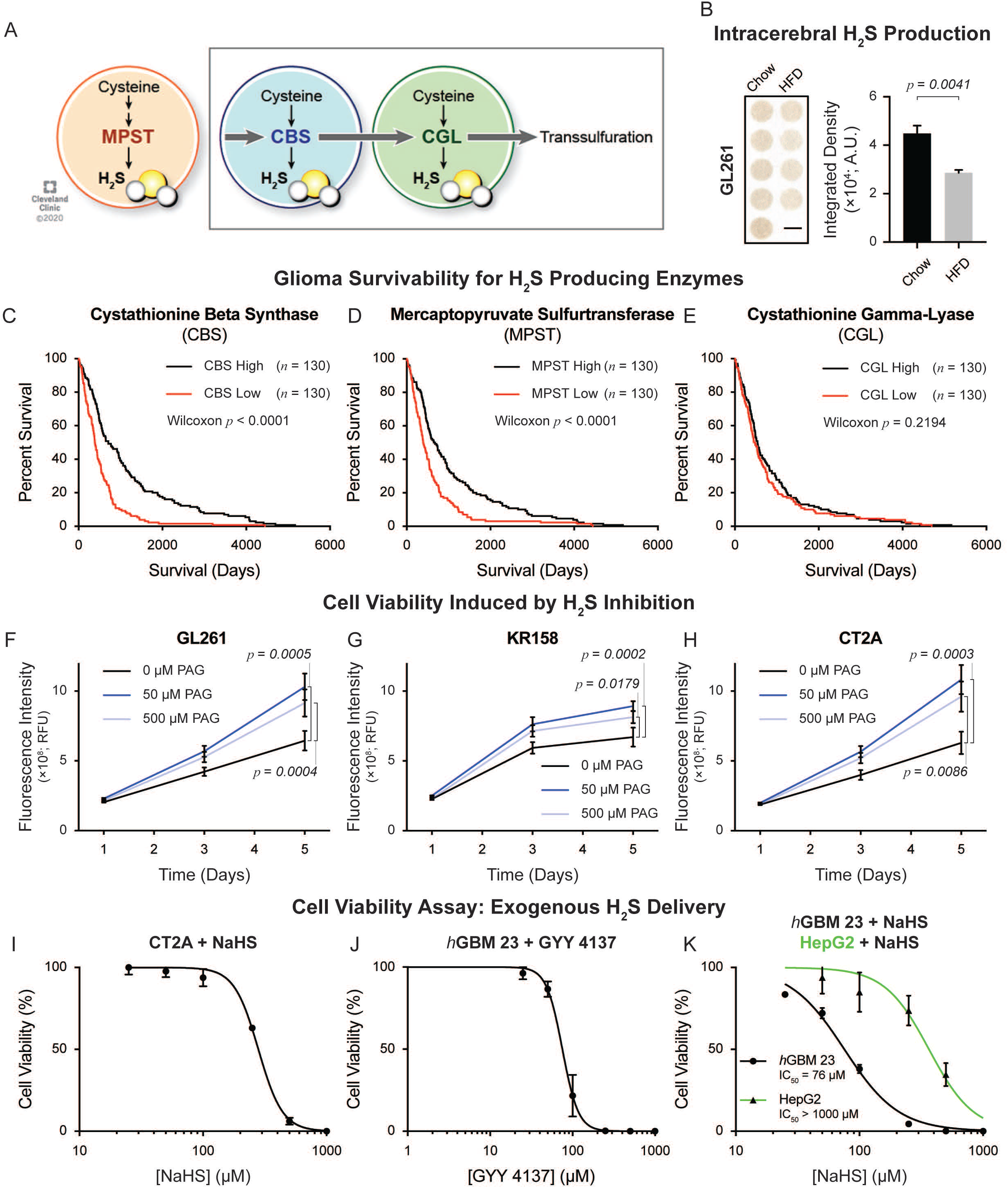
HFD and Gliomagenesis Inhibit Hydrogen Sulfide Production, Resulting in Tumor Cell Hyperproliferation. (**A**) Schematic detailing the generation of H_2_S as a byproduct of cysteine metabolism associated with MPST activity and the transsulfuration pathway. (**B**) H_2_S production analysis indicates that HFD consumption results in decreased H_2_S synthesis in the tumors of HFD-fed mice as compared to tumors isolated from mice fed a control diet. Each well contains brain and tumor tissue homogenate from separate and distinct experimental animals. *p* values determined by unpaired t-test. (**C, D**) Human patient data curated by The Cancer Genome Atlas Low-Grade Glioma and GBM dataset indicate that diminished mRNA expression of the H_2_S-generating enzymes CBS and MPST predicts poor overall survival. (**E**) While the mRNA expression of CGL did not directly inform patient survival, previously published biochemical analysis confirmed that this enzyme is non-functional in the context of GBM. (**F – H**) Cell-Titer Glo viability analysis confirmed that *in vitro* treatment with the CGL-selective inhibitor propargylglycine (PAG) increased GL261, KR158, and CT2A tumor cell viability compared to vehicle controls. *p* value determined by two-way ANOVA; 5 technical replicates per timepoint. (**I – K**) H_2_S supplementation using the chemical donor sodium hydrosulfide (NaHS) or GYY4137 resulted in selective viability reduction for mouse (CT2A) and human (*h*GBM 23) tumor cells compared to non-GBM (HepG2) cellular controls. IC_50_ values were determined based on nonlinear regression analysis. Five technical replicates were analyzed per concentration. While representative IC_50_ curves for CT2A, *h*GBM 23 and HepG2 are depicted here, IC_50_ concentrations were determined for a total of 3 human GBM specimens, 2 syngeneic GBM specimens, and 2 non-GBM cellular controls.

### HFD Consumption and Not Obesity Drives GBM Acceleration

There are a number of cancers, including hepatocellular carcinoma (HCC), that are accelerated by HFD consumption but also by the metabolic state that accompanies obesity regardless of diet*(31*, *32*, *40)*. Our experimental animals were primed with HFD for 2 weeks prior to tumor introduction; however, their body fat percentage did not reflect obesity (≥ 25%) until much later (~ 2 weeks) in the experiment (**Supplementary Figure 1B**). Because obesity, and not diet, is the predominant clinical variable that is collected and used as an epidemiological benchmark, we wanted to understand the degree to which obesity contributed to GBM acceleration separate from HFD consumption. We therefore turned to the LepOB mutant mouse, which exhibits many of the hallmark physiological features of metabolic syndrome, including obesity (**Supplementary Figure 4A, B**), as a result of hyperphagic consumption of standard rodent chow. We compared the overall survival of tumor-bearing LepOB mice and C57BL/6J mice, both fed a control, low-fat diet. Under conditions of obesity but in the absence of HFD, no GBM acceleration was observed. Overall survival was not significantly different between tumor-bearing LepOB and C57BL/6J mice (**Supplementary Figure 4C**). Further, analysis of the excised endpoint tumors revealed identical capacities for H_2_S production (**Supplementary Figure 4D**). Thus, H_2_S synthesis inhibition, which accelerated GBM progression, required HFD consumption. The metabolic profile associated with obesity was not sufficient to drive hyper-aggression in GBM.

### Enhanced Metabolic Substrate Fluidity Results from H_2_S Inhibition

To this point, we focused exclusively on experimental animals held under precisely controlled dietary conditions for extended periods of time. The degree to which H_2_S tumor suppression translates to the human condition remains a critical and unresolved question. We collected and analyzed tissues from 5 patients and 5 non-cancerous control brain specimens that had been flash frozen at the time of isolation. Initial examination for the ability to produce H_2_S revealed that GBM tissue produced approximately 50% of this critical tumor suppressor compared to non-cancerous control tissues (**Figure 4A**). It is worth noting here that we experimentally attenuated H_2_S production to a similar degree utilizing HFD (**Figure 3B**). Leveraging a mass spectrometry-based method, we further analyzed these specimens to generate a complete profile of the S-sulfhydrated proteins contained within. Consistent with the reduced H_2_S production, we noted a significant decrease in the number of S-sulfhydrated proteins within the GBM specimens compared to controls (**Figure 4B**). S-sulfhydration loss affected more than 161 separate proteins mechanistically involved in multiple molecular pathways. We then stratified the S-sulfhydrated-protein landscape into biochemical pathways using KEGG Pathway analysis (**Figure 4C**). We were struck by the number of metabolic pathways represented in this analysis. In the context of GBM, protein S-sulfhydration was dysregulated across multiple metabolic pathways. Carbon metabolism, pyruvate and amino acid metabolism, oxidative phosphorylation, and glycolysis were all significantly impacted through the loss of H_2_S signaling. This degree of metabolic reprogramming likely enables the substrate flexibility and enhanced adaptability that define the GBM CSC phenotype and helps explain the hyper-aggressive disease presentation brought about by HFD consumption in mice. These data indicate that S-sulfhydration loss results in a broad-spectrum molecular reprogramming that enables dynamic metabolic adaptability within GBM tumors.

**Figure 4.**
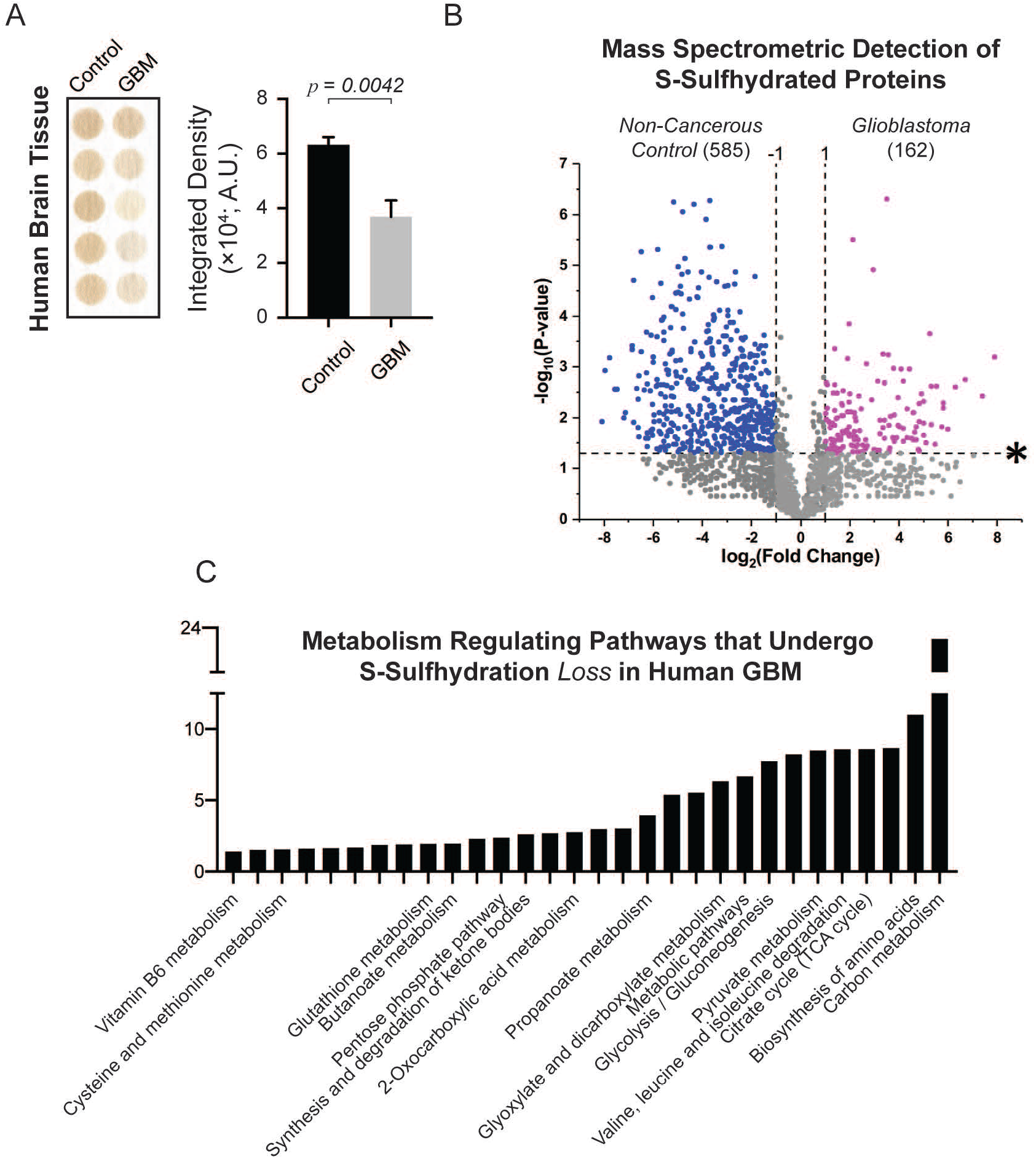
Gliomagenesis Induces Significant Loss in H_2_S Synthesis and Signaling Primarily Associated with Cellular Metabolism. (**A**) H_2_S production analysis confirms that human GBM tumors produce less H_2_S than non-cancerous control brain tissues. Each well contains brain or tumor tissue homogenate from separate biopsy specimens. *p* values determined by unpaired t-test. (**B**) Volcano plot representing the LC-MS S-sulfhydration analysis reveals striking deficits in the post-translational H_2_S signaling profile of human GBM as compared to non-cancerous human brain tissue. (**C**) KEGG pathway analysis of the proteins that have undergone S-sulfhydration *loss* in the context of GBM identifies a broad-spectrum molecular reprogramming centered on GBM tumor cell metabolism.

## Discussion

These findings confirm that HFD consumption accelerates and intensifies GBM in the preclinical experimental setting. The degree to which this finding extends to the human clinical condition remains an open question. As long-term dietary patterns are not accounted for as a variable in most clinical settings, epidemiological studies primarily question whether obesity, as measured by body mass index (BMI), represents a risk factor for GBM initiation. Such a link has been demonstrated for several cancers*(31*, *32)*; however, the connection between gliomagenesis and obesity has been inconsistent. For example, the recent completion of a massive meta-analysis leveraging clinical data from more than 10 million subjects led to the conclusion that overweight (BMI: 25 – 30 ^kg^/_m_^2^) and obese (BMI > 30 ^kg^/_m_^2^) status represents a risk factor for glioma development specifically for women*(41)*. However, other reports fail to substantiate this finding*(42*, *43)*. Little to no attention has been paid to the possibility that a long-term pattern of HFD consumption may profoundly alter GBM evolution as it develops within the brain. Based on these findings, it is conceivable that patients who consume a HFD long-term could experience a hyper-aggressive disease trajectory and/or present disease that is more adaptable and therefore more challenging to treat. Our findings on HFD-induced adult neurogenesis (**Supplementary Figure 2**) suggest that HFD consumption may facilitate GBM development by expanding the NSPC population, which has been identified as a potential cell of origin for GBM*(44*, *45)*. Additionally, our *in vivo* limiting-dilution experiments (**Figure 1E – G**) indicated that GBM may be better able to take root and thrive in the HFD-fed brain. Recently, a diet-focused meta-analysis of 1.2 million subjects was presented involving self-reported dietary patterns. Once again, investigators were limited to interrogating this dataset for risk factor identification, concluding that no specific diets served as risk factors for GBM initiation*(43)*. Including factors such as progression-free survival and long-term dietary pattern will help to clarify the degree to which obesogenic diets modulate the course of human GBM. Based on that additional level of understanding, we will be better able to evaluate diet as a prognostic indicator and manage our expectations for using diet as a disease management tool.

There are interesting overlaps between this study and work on the ketogenic diet as an anti-GBM intervention. Both lines of research embrace the idea that diet might have profound effects on the trajectory of disease. The ketogenic diet, which is commonly employed against epilepsy, is typically formulated as an extremely high-fat (90%), low-carbohydrate (5%) diet. Recently, it has been hypothesized that the glucose deprivation that results from strict adherence to the ketogenic diet might enable long-term GBM management by starving these tumors of their preferred metabolic substrate*(29*, *46)*. Unfortunately, a synthesis of recent work in GBM metabolism confirms that these tumors are not strictly dependent on glucose. Instead, subpopulations of tumor cells present a propensity for metabolic adaptability. A recent study of glucose starvation in GBM confirmed that while the bulk of the malignant population, made up of non-stem tumor cells, were clearly dependent on glucose, CSCs were metabolically plastic. They were uniquely capable of adapting to multiple metabolic substrates*(11*–*13*, *47)*, especially favoring fatty acid oxidation in the context of glucose starvation. While this strategy effectively eliminated the glucose-dependent non-stem tumor cell population, GBM CSCs bypassed this metabolic dependency and expanded into the treatment-induced vacancy*(14)*. In agreement, we found that HFD consumption altered the nutrient landscape of the brain, specifically resulting in intracerebral accumulation of saturated lipids. As seen in the liver, this lipid excess inhibited H_2_S production and signaling, in turn driving CSC enrichment and disease acceleration. These findings caution against the use of the ketogenic diet as a long-term GBM management tool. The ketogenic diet represents a selective pressure, similar to a targeted chemo-, radio-, or immunotherapy. The inherent heterogeneity, complexity, and adaptability of GBM will ultimately drive the evolving tumor to sidestep any singular selective pressure.

Our findings also highlight a post-translational modification that is profoundly altered in GBM and has received very little attention. H_2_S and S-sulfhydration have been a focus of research in the aging*(48*, *49)*, neurodegeneration*(21)*, and metabolism fields; however, there is very little known about this metabolite in GBM tumors*(26*, *27)*. Our work reinforces the concept of H_2_S signaling as a tumor suppressor for GBM while simultaneously introducing numerous questions about the mechanisms by which this is achieved. For example, *what is the mechanism responsible for H_2_S signaling attenuation in GBM*? Previous work established that attenuation of synthesis resulted from down-regulation of CGL, CBS, and MPST. HFD consumption can decrease the expression of these enzymes at the protein level, as was demonstrated previously in the livers of HFD-fed wild-type animals*(20)*. Synthesis attenuation can also result from enzyme loss of function, which has been confirmed in the context of human GBM*(27)*. Additionally, *by what mechanism does S-sulfhydration loss drive GBM progression*? Here, it is worth noting that the functional consequences of S-sulfhydration have only been reported for a limited number of proteins. In a recent review, the functional changes that resulted from S-sulfhydration were highlighted for 43 cysteine residues present within only 25 total proteins*(22)*. Thus, at this point, we can only speculate based on our S-sulfhydration analysis that the loss of H_2_S signaling serves to increase the tumor’s capacity for metabolic adaptability and mitochondrial fitness. A preliminary *in silico* examination revealed three metabolism-associated proteins: fatty acid-binding protein 3 (FABP3), enoyl-CoA hydratase 1 (ECH1), and ATP synthase peripheral stalk subunit OSCP (ATP5O), which established a presumptive signaling pathway that can be tested in the future for its capacity to increase metabolic fitness and fatty acid utilization based on the loss of S-sulfhydration (**Table 2**). If loss of S-sulfhydration on these three proteins conferred enzymatic gain of function, then FABP3 would more effectively channel fatty acids into the cytosol of GBM tumor cells, ECH1 would enhance the lipid β-oxidation cycle, and ATP5O may increase a cell’s capacity for oxidative phosphorylation. Gain of function for these three proteins is consistent with functional changes that result from S-sulfhydration of factors that regulate metabolism and mitochondrial fitness *(22)*. Lastly, *could this non-traditional, non-genetic tumor suppressor be replaced or supplemented in conjunction with standard of care to better manage GBM in human patients*? There are a variety of chemical H_2_S donors, pharmaceutical H_2_S inducers, and diets that drive endogenous H_2_S production. A major challenge here will be one of bio-availability. For an H_2_S replacement strategy to work, the metabolite must be relatively stable and able to penetrate the brain. While we are encouraged by our *in vitro* H_2_S treatment data (**Figure 3I – K**), currently available donors and inducers fall short of these standards *in vivo*. Nonetheless, we suggest that pursuing strategies to induce intracerebral H_2_S production and signaling may serve to limit GBM metabolic adaptability, making the disease a more receptive target for cytotoxic therapies.

In conclusion, we have demonstrated that consumption of an obesogenic HFD resulted in a shift in the nutrient landscape of the brain. The resulting attenuation of tumor-suppressive and metabolism-suppressive H_2_S enabled adaptation to this lipid excess. These fundamental changes within the brain and tumor microenvironment induced CSC enrichment, heightened chemotherapy resistance, and accelerated GBM progression (**Figure 5**).

**Figure 5.**
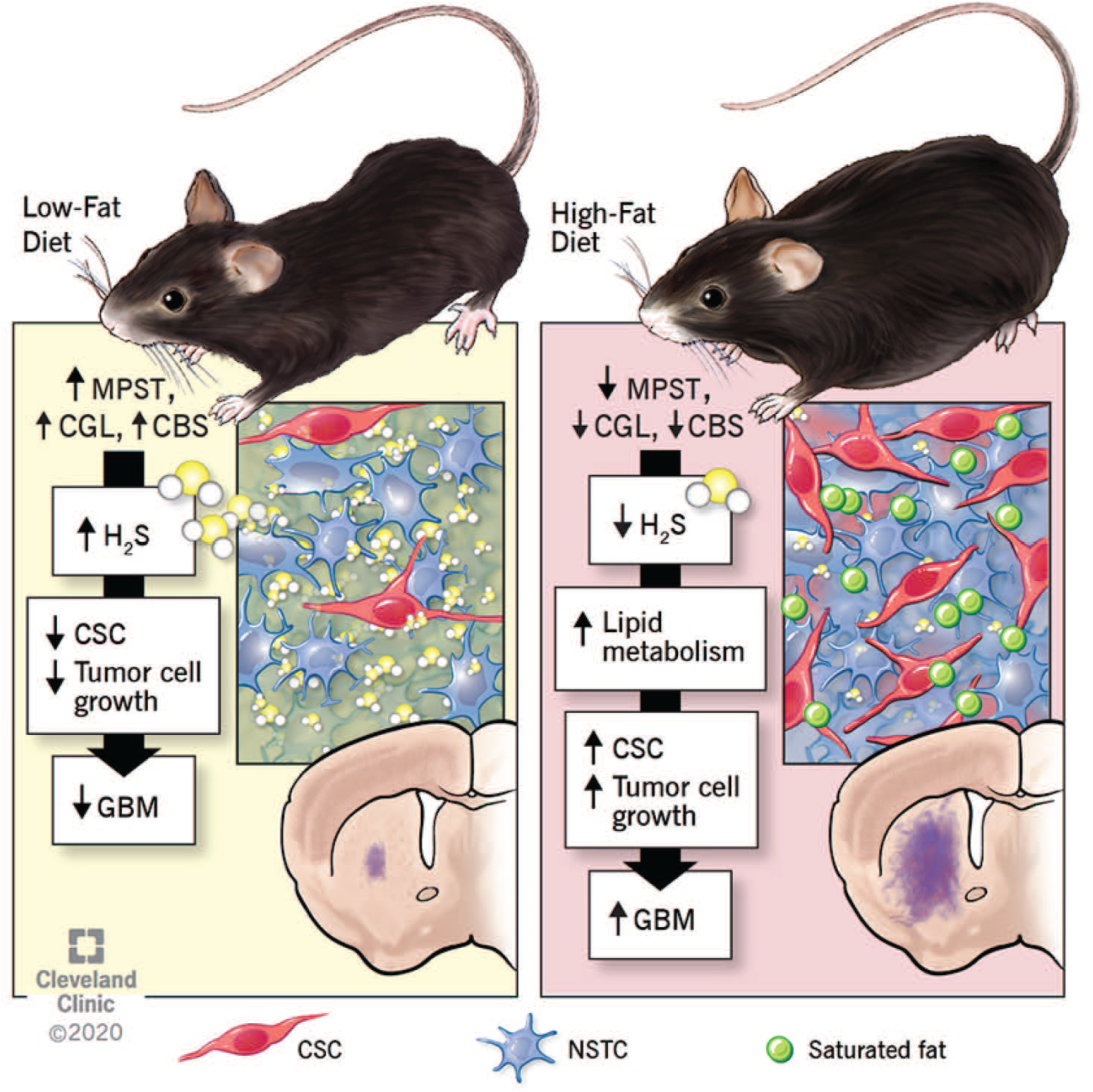
Consumption of HFD Inhibits the Tumor-Suppressive Activity of H_2_S, Driving CSC Enrichment and Progression in GBM. In total, our findings indicate that HFD consumption serves as an accelerant for GBM. HFD consumption drives accumulation of saturated, mono-unsaturated and di-unsaturated fats within the brain. This lipid excess inhibits H_2_S production, which results in a wide-ranging attenuation of S-sulfhydration centered on tumor metabolism regulators. These linked intrinsic and extrinsic changes within the tumor result in the expansion of treatment-refractory CSCs within the tumor microenvironment and truncated overall survival.

## Materials and Methods

For detailed purchasing and use instructions, please refer to **Table 3**.

### Cell Culture

The patient-derived GBM models (*h*GBM 23, *h*GBM, 124, *h*GBM 3832, *h*GBM 3691) as well as the syngeneic mouse GBM models (CT2A and KR158) were propagated under adherent culture conditions. Single-cell suspensions of 50,000 cells/mL were suspended in DMEM/F12 media enriched with N2 Supplement, *rh*EGF, *rh*FGF, and penicillin-streptomycin and seeded into Geltrex-coated culture flasks.

The syngeneic mouse GBM model GL261 and the liver cancer control models HepG2 and NCTC were propagated under adherent culture conditions in DMEM/F12 supplemented with 10% fetal bovine serum (FBS) and penicillin-streptomycin.

For all cellular models, complete media including supplements were exchanged every other day until the cultures reached 85 – 90% confluence. Cultures were passaged weekly using Accutase. Phosphate-buffered saline was used to wash, quench, and collect cells before re-plating. Cells were grown and maintained in humidified incubators held at 37°C and 5% CO_2_.

The patient-derived GBM models *h*GBM 23 and *h*GBM 124 were obtained from Dr. Erik Sulman (New York University). The patient-derived GBM model *h*GBM 3691 was obtained from Dr. Jeremy Rich (University of California San Diego). The syngeneic GBM models CT2A and KR158 were obtained from Dr. Loic Deleyrolle (University of Florida).

### Differential Diet and Intracerebral Tumor Cell Implantation

All animal procedures were evaluated and approved prior to initiation by the Institutional Animal Care and Use Committee (IACUC) of the Cleveland Clinic Lerner Research Institute. For comparative survival studies concerning high-fat *versus* control diets, 4-week-old female mice were purchased from Jackson Laboratories. Based on the patient epidemiological data, which indicated that overweight and obese status represents a significant risk for gliomagenesis in women only, we elected to perform these *in vivo* studies exclusively in female mice. Mice were subdivided into two groups: (1) a HFD-fed group and (2) a control diet-fed group. For the syngeneic, mouse models, C57BL/6J mice were used, and the control group was fed standard rodent chow. For the patient-derived GBM model, NOD-SCID mice were employed, and the control group was fed a low-fat, energy-balanced diet. Differential diets were introduced 2 weeks prior to tumor cell implantation and maintained throughout the post-implantation survival period. Diet was completely exchanged once per week to prevent spoilage. For tumor cell implantation, 6-week-old, diet-primed mice were anesthetized using inhaled isoflurane (2–2.5%) and fit to a stereotaxic apparatus. Using an insulin syringe secured to a large probe holder, the 31-guage needle was passed directly through the scalp and skull approximately 0.5 mm rostral and 1.8 mm lateral to bregma. The needle was lowered 3.0 mm beneath the surface of the scalp, where a specific number of cells (50,000; 20,000; 15,000; 10,000; 5,000) suspended in 5 μL of DMEM/F12 were slowly injected. The needle was held in place an additional 60 seconds before a slow and measured removal. Animals were monitored over time for changes in body mass, fat-to-lean mass composition using EchoMRI, and the presentation of the set of neurological and behavioral symptoms associated with end-stage brain cancer (**Figure 1A**).

### Tissue Preparation and Immunofluorescence Analysis

At the experimental endpoint, animals were subdivided into one of two possible tissue preservation and harvesting modalities. Approximately 30% of endpoint animals were anesthetized and subjected to cardiac perfusion with 4% formaldehyde in PBS. The brain was removed and post-fixed overnight. The tissue was then cryoprotected with sequential treatments first in a solution of 30% sucrose in PBS and then in a 1:1 mixture of 30% sucrose (PBS):O.C.T. Compound. Finally, the tissue was embedded in O.C.T Compound, and 20 μm coronal sections were prepared for subsequent immunofluorescence analysis using standard protocols. The specific antibodies used can be found in **Table 1**. In all cases, primary antibodies were coupled to appropriate Alexa Fluor 488- or 555-conjugated secondary antibodies. Engrafted syngeneic tumor cells were distinguished from the surrounding host brain tissue based on the expression of MCM2. The cancer stem cell phenotype was evaluated based on expression of the transcription factor SOX2, and nuclei were visualized with Hoechst 33342. The stained sections were mounted onto slides, coverslipped, and examined using an inverted Leica SP8 confocal microscope.

The remaining 70% of endpoint animals were anesthetized and subjected to cardiac perfusion with PBS. The brain was dissected away from the calvarium and bisected along the midline, generating a pair of matched samples: (1) The tumor-bearing hemisphere and (2) the contralateral, healthy control hemisphere. Labeled samples were immediately flash frozen in liquid nitrogen and stored at −20°C until downstream tissue analysis.

### Untargeted Mass Spectrometry-Based Lipidomic Analysis

To characterize the lipid landscape of the HFD- *versus* chow-fed brain and tumor, we employed a mass spectrometry-based shotgun lipidomic analysis method that enables identification of multiple structurally distinct lipid species. This method, which we adopted without alteration, was originally published by Gromovsky *et al*. *(50)* and later optimized by Neumann *et al*. *(51)*.

### Preparation of Oleic and Linoleic Acid

The free fatty acids oleic and linoleic acid were delivered to cultured cells bound to BSA. Therefore, we initially generated a 3% (m/v) fatty-acid free bovine serum albumin solution using D-PBS at room temperature. Oleic acid and linoleic acid were solubilized separately in 90% aqueous ethanol, producing a 50 mM free fatty acid (FFAs) solution. These solutions were then diluted to 50 nM stock solutions in the 3% BSA solution, aliquoted, and stored at −20°C until needed.

### Tumor Cell Proliferation Assessment

#### 1. Cell-Titer Glo Viability Assay

Proliferation was assessed in the contexts of excess oleic and linoleic acid, in the presences of PAG, and in response to the exogenous H_2_S donors NaHS and GYY4137 using the Cell-Titer Glo viability assay following the protocol established by the manufacturer. A total of 2000 cells/well were plated onto Geltrex-coated 96-well plates. BSA-bound oleic and linoleic acids were added to the media at the following concentrations: Vehicle (3% BSA alone), 100 nM, 1 μM, and 10 μM. All conditions were plated in 5 technical replicates. Readings were taken on the day of plating (Day 0) as well as on Days 1, 3, and 5. Cell viability was normalized to Day 0 to account for any plating irregularity.

#### 2. Direct Cell Counting

Cellular proliferation was validated under conditions of H_2_S inhibition through direct cell counting. In this case, 20,000 cells/well were plated onto Geltrex-coated 12-well plates. Vehicle (DMSO), PAG (solubilized in growth media), or temozolomide (TMZ, solubilized in DMSO) was added at the time of plating. All conditions were plated in four technical replicates. Cells were dissociated, collected, and counted with an automated hemocytometer 3 days after plating.

### In Vitro Limiting-Dilution Analysis (LDA)

Extreme LDA was performed to assess self-renewal in the context of excess mono- and di-unsaturated lipids. Cells were diluted progressively throughout the plate beginning with 100 cells/well across the first row. Subsequent dilutions resulted in the delivery of 50, 25, 13, 6, 3, 1, and 0 cells/well. Separate plates were prepared containing vehicle (3% BSA Alone), 10 μM, 50 μM, and 100 μM oleic or linoleic acid bound to 3% BSA. Cells were maintained in culture for 10 – 14 days. Fifty microliters of media + lipids or media + vehicle was added to the appropriate wells every 5-7 days. Each well was scored as either positive or negative for sphere outgrowth. The Walter and Eliza Hall Institute Bioinformatics Division ELDA analyzer*(52)* was used to analyze data and calculate stem cell frequency.

### IC_50_ Assessment of the H_2_S Donors NaHS and GYY4137

The half maximal inhibitory concentration (IC_50_) was established for the H_2_S donors NaHS and GYY4137 across a variety of cellular models. Fresh NaHS was solubilized in D-PBS at the time of experimental setup. GYY4137 was solubilized in 100% ethanol and stored at −20°C. A total of 1000 cells/well was plated in white-walled, Geltrex-coated 96-well plates. Five technical replicates were plated per H_2_S donor concentration. Wells were then supplemented with either NaHS or GYY4137 at the following concentrations: 10 μM, 25 μM, 50 μM, 75 μM, 100 μM, and 500 μM along with appropriate vehicle controls. Following the manufacturer’s instructions, cellular viability was determined using the Cell-Titre Glo assay after 5 days of incubation.

### Lead Acetate/Lead Sulfide H_2_S Production Assay

The endogenous H_2_S production capacity of tissues (and pelleted cells) was measured using the lead acetate/lead sulfide method. This method, which we adopted without alteration, was originally published by Hine and colleagues*(48)*. H_2_S production was assessed from tumor-containing specimens collected from HFD- *versus* control-fed mice, LepOB *versus* C57BL/6J mice, and human GBM *versus* non-cancerous control specimens. Age-matched and sex-matched GBM and non-cancerous control brain specimens were obtained from the Cleveland Clinic Burkhardt Brain Tumor Center under IRB 2559.

### Protein S-Sulfhydration Landscape Analysis

To characterize the S-sulfhydration landscape of human GBM tumors and non-cancerous control tissues, we adopted a multi-stage method adapted and optimized by our collaborators. This method, which we employed without alteration, was recently described by Bithi and colleagues*(53)*.

### Statistical Analyses for S-Sulfhydration Profiling

Microsoft Excel, GraphPad Prism and OriginPro were used for data analysis and statistics. Microsoft Excel and GraphPad Prism data are presented here as mean ± SEM. The difference between two groups was analyzed by unpaired t-test, with the significance level set to *p* < 0.05. OriginPro was used to generate volcano plots in which the X-axis reported the fold change between GBM and non-cancerous control groups represented in log_2_ scale. The Y-axis reported negative log_10_ of the p-values from the comparative t-tests.

### Statistical Analyses for In Vitro and In Vivo Experimentation

All experiments were performed in triplicate. Results are reported mean ± SEM. Unless otherwise stated, one-way ANOVA was used to calculate statistical significance; *p*-values and sample size (*n*) are detailed in the text and figure legends. *In vivo* survival analysis was calculated by log-rank analysis.

### Interrogation of The Cancer Genome Atlas Database

Gene expression data were obtained from the University of California Santa Cruz Xena Functional Genomics Explorer*(54)*, leveraging data from The Cancer Genome Atlas Low-Grade Glioma and Glioblastoma study. Expression levels were categorized into two groups based on median gene expression. Log-rank survival analysis was performed according to the overall survival times and status for patients in both groups.

## Acknowledgements

We would like to thank the members of the Lathia, Hine, and Brown laboratories for insightful conversations and constructive criticism. We also thank Dr. Belinda Willard of the Lerner Proteomics and Metabolomics Core for assistance with mass spectrometry, Amanda Mendelsohn from the Cleveland Clinic Center for Medical Art & Photography for illustration assistance. We thank Drs. Erik Sulman, Loic Deleyrolle, and Jeremy Rich for providing syngeneic and patient-derived GBM models. This work was supported by pilot funding from the Case Comprehensive Cancer Center (C.H., J.D.L., D.J.S). The Lathia Laboratory is supported by the National Institutes of Health (R01 NS089641, R01 NS109742, and R01 NS083629), a Sontag Foundation Distinguished Scientist Award, and the VeloSano Bike Race. The Hine Laboratory is supported by the National Institutes of Health (R01 HL148352 and R00 AG050777). The Brown Laboratory receives funding from the National Institutes of Health (R01 DK120679 and P50 AA024333). D.J.S. was supported by an NIH Kirschstein NRSA (F32CA213727).

---

**Supplementary Figure 1.**
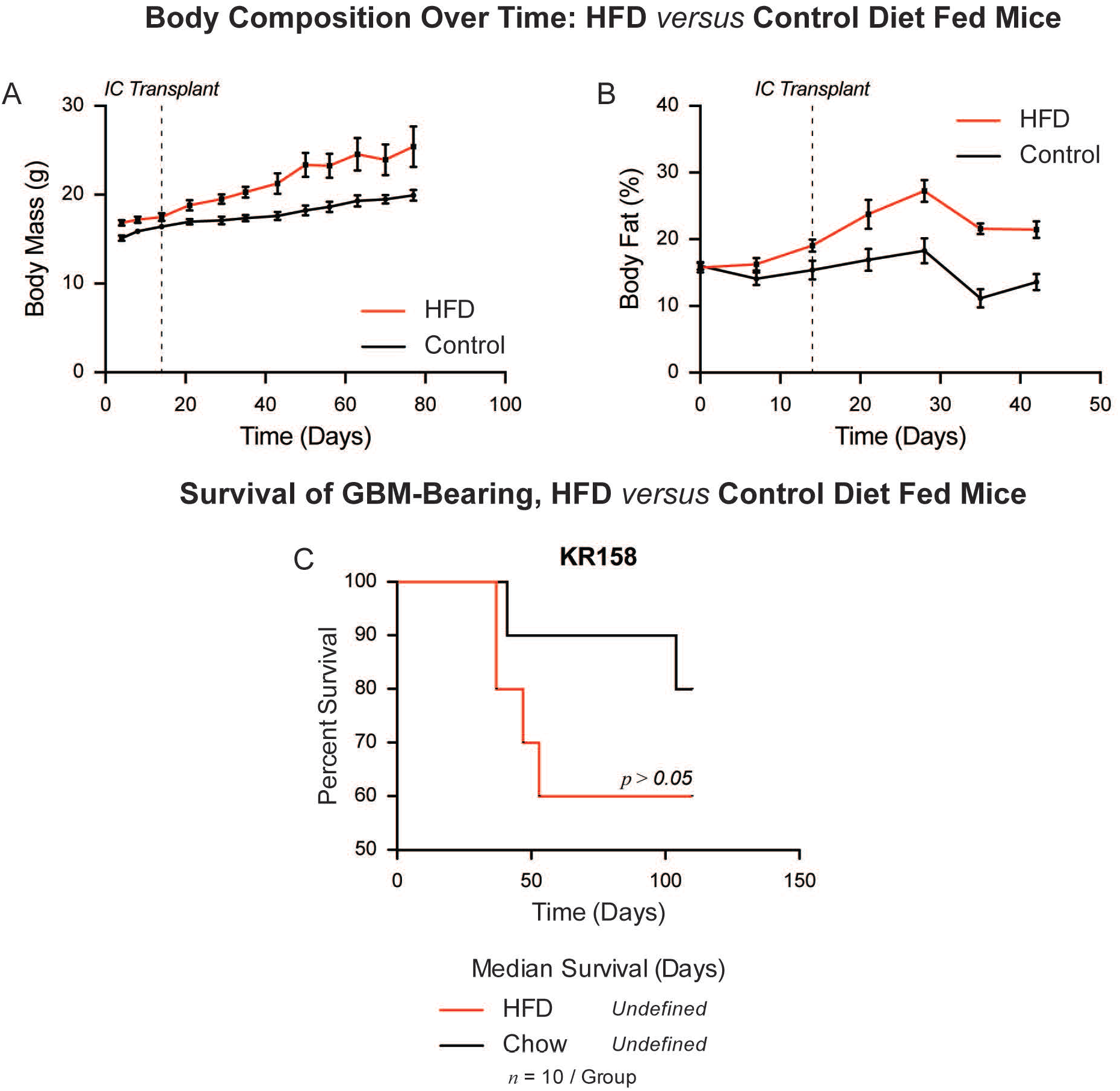
Obesogenic HFD Consumption Leads to Body Fat Accumulation Over Time. (**A, B**) Representative assessments of body mass and fat compositional changes over time as a function of HFD *versus* control diet consumption. Each *in vivo* experiment was tracked using these metrics beginning from the introduction of the differential diets throughout the post-transplantation period. (**C**) For any cell dosage, Kaplan-Meier survival analysis of the syngeneic GBM model KR158 demonstrated the expected pattern of HFD-mediated disease acceleration; however, statistical testing was confounded based on imperfect disease penetrance.

**Supplementary Figure 2.**
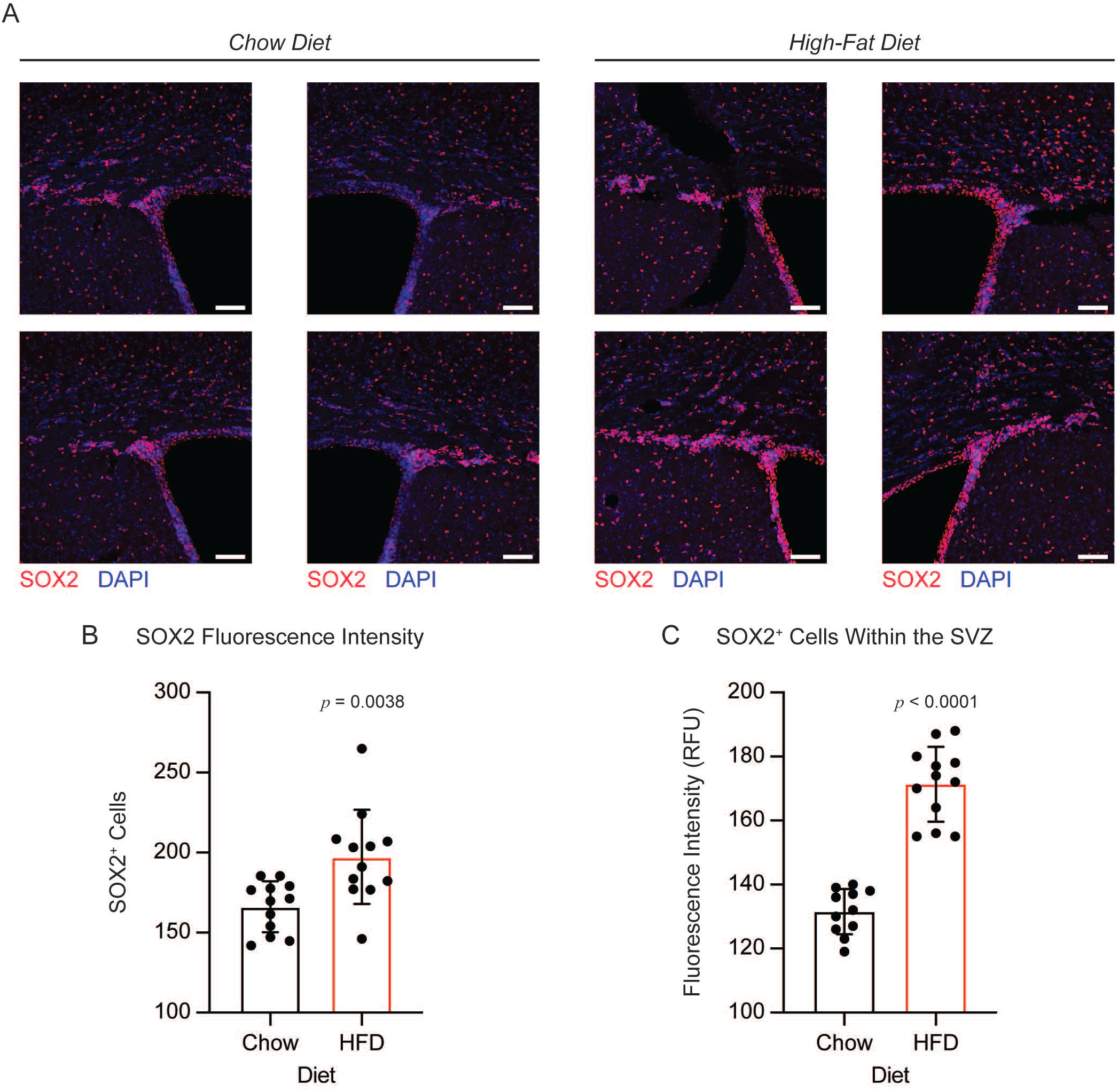
The HFD-Fed Brain Fosters Expansion of the Neural Stem and Progenitor Cell Pool within the Subventricular Zone. (**A**) Representative bilateral immunofluorescent micrographs of the subventricular germinal zone of HFD- *versus* chow-fed mice. The neural stem cell-associated transcription factor SOX2 was visualized in red, and nuclei were visualized in blue using DAPI. A total of 12 representative images were captured from a total of 3 animals per group. Quantification based on (**B**) SOX2 fluorescence intensity or (**C**) direct counting of nucleated SOX2^+^ cells confirmed expansion of the NSPC population under HFD-fed conditions as compared to control diet. *p* values determined by unpaired t-test.

**Supplementary Figure 3.**
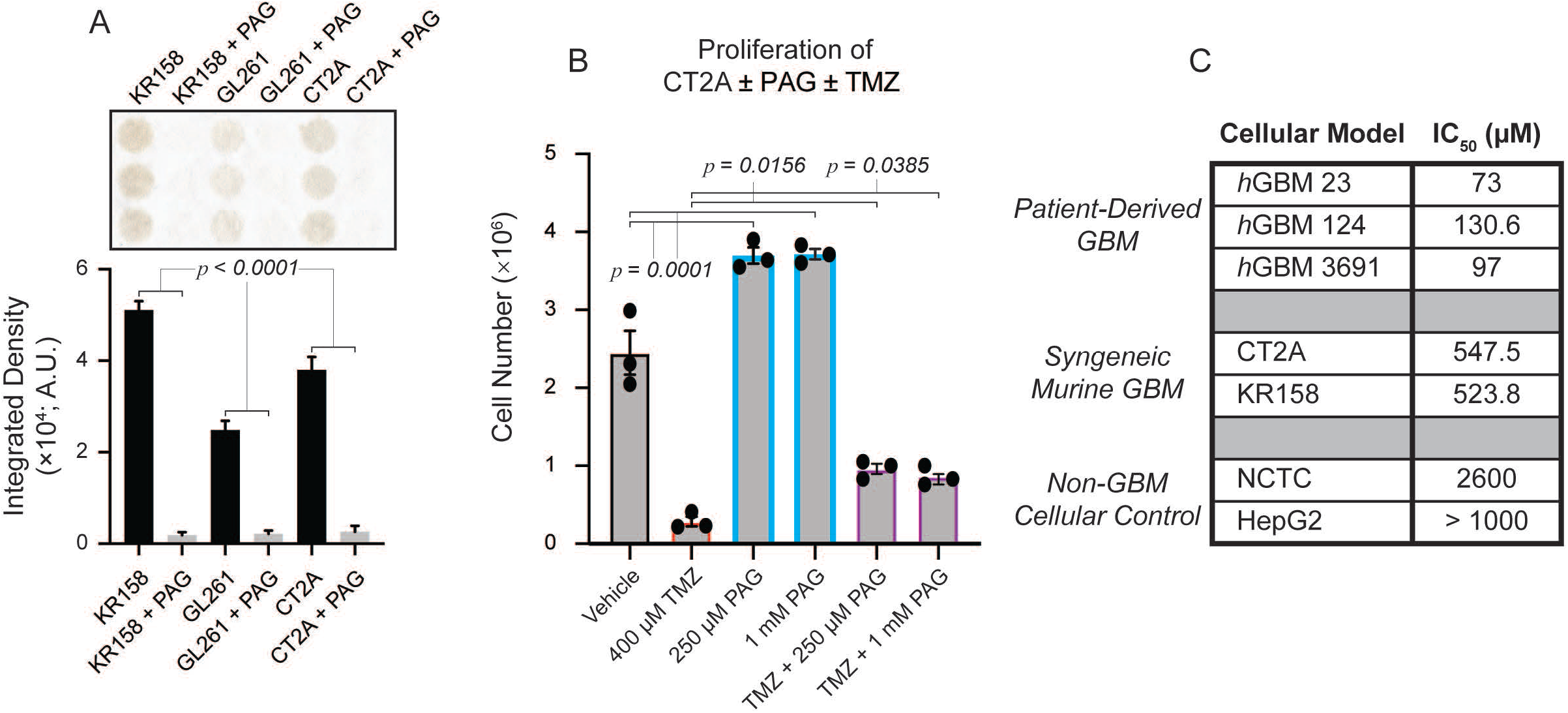
Inhibition of H_2_S Synthesis Drives Tumor Cell Proliferation and Chemotherapy Resistance. (**A**) *In vitro* verification that the CGL inhibitor PAG effectively attenuates H_2_S production from the syngeneic GBM models KR158, GL261, and CT2A. Each well contains lysate from separate biological replicates. For a given cellular model, *p* value was determined using an unpaired t-test. (**B**) Direct cell counting verifies that H_2_S inhibition increases tumor cell proliferation and resistance to the standard-of-care chemotherapeutic temozolomide (TMZ).

**Supplementary Figure 4.**
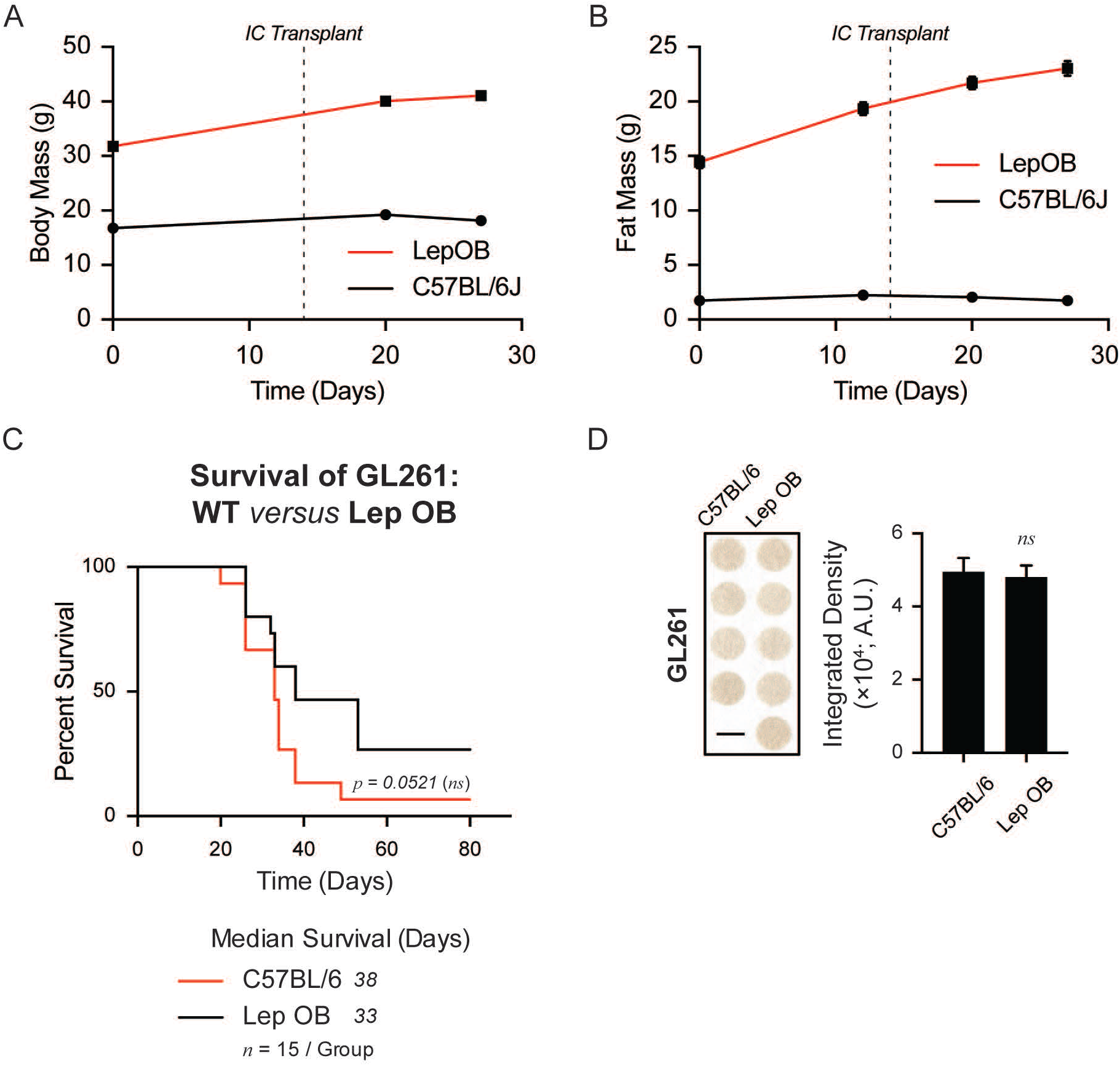
HFD Consumption Rather than Genetic Obesity Accelerates GBM and Attenuates H_2_S Synthesis. (**A, B**) Representative assessments of body mass and fat compositional changes over time in the context of leptin knockout *versus* wild-type C57BL/6J genetics. (**C**) Kaplan-Meier survival analysis indicates that overall survival between GBM-bearing C57BL/6J and genetically obese LepOB mice was mathematically indistinguishable as determined by log-rank statistical analysis. Additionally, (**D**) intracerebral H_2_S synthesis remained intact in the context of genetic leptin knockout. Each well contains brain and tumor tissue homogenate from separate and distinct experimental animals. *p* values determined by unpaired t-test.

## References

1. B. Tran, M. A. Rosenthal, Survival comparison between glioblastoma multiforme and other incurable cancers, Journal of Clinical Neuroscience 17, 417–421 (2010).

2. M. Weller, W. Wick, K. Aldape, M. Brada, M. Berger, S. M. Pfister, R. Nishikawa, M. Rosenthal, P. Y. Wen, R. Stupp, G. Reifenberger, Glioma, Nat. Rev. Dis. Primers 39, 15017–18 (2015).

3. A. Sottoriva, I. Spiteri, S. G. M. Piccirillo, A. Touloumis,V. P. Collins, J. C. Marioni, C. Curtis, C. Watts, S. Tavaré, Intratumor heterogeneity in human glioblastoma reflects cancer evolutionary dynamics, Proceedings of the National Academy of Sciences 110, 4009–4014 (2013).

4. A. P. Patel, I. Tirosh, J. J. Trombetta, A. K. Shalek, S. M. Gillespie, H. Wakimoto, D. P. Cahill, B. V. Nahed, W. T. Curry, R. L. Martuza, D. N. Louis, O. Rozenblatt-Rosen, M. L. Suvà, A. Regev, B. E. Bernstein, Single-cell RNA-seq highlights intratumoral heterogeneity in primary glioblastoma, Science 344, 1396–1401 (2014).

5. R. Reinartz, S. Wang, S. Kebir, D. J. Silver, A. Wieland, T. Zheng, M. Küpper, L. Rauschenbach, R. Fimmers, T. M. Shepherd, D. Trageser, A. Till, N. Schäfer, M. Glas, A. M. Hillmer, S. Cichon, A. A. Smith, T. Pietsch, Y. Liu, B. A. Reynolds, A. Yachnis, D. W. Pincus, M. Simon, O. Brüstle, D. A. Steindler, B. Scheffler, Functional Subclone Profiling for Prediction of Treatment-Induced Intratumor Population Shifts and Discovery of Rational Drug Combinations in Human Glioblastoma, Clin Cancer Res 23, 562–574 (2016).

6. F. A. Siebzehnrubl, D. J. Silver, B. Tugertimur, L. P. Deleyrolle, D. Siebzehnrubl, M. R. Sarkisian, K. G. Devers, A. T. Yachnis, M. D. Kupper, D. Neal, N. H. Nabilsi, M. P. Kladde, O. Suslov, S. Brabletz, T. Brabletz, B. A. Reynolds, D. A. Steindler, The ZEB1 pathway links glioblastoma initiation, invasion and chemoresistance, EMBO Mol Med, n/a–n/a (2013).

7. J. D. Lathia, J. Gallagher, J. M. Heddleston, J. Wang, C. E. Eyler, J. Macswords, Q. Wu, A. Vasanji, R. E. Mclendon, A. B. Hjelmeland, J. N. Rich, Integrin alpha 6 regulates glioblastoma stem cells, Cell Stem Cell 6, 421–432 (2010).

8. T. L. Haas, M. R. Sciuto, L. Brunetto, C. Valvo, M. Signore, M. E. Fiori, S. di Martino, S. Giannetti, L. Morgante, A. Boe, M. Patrizii, U. Warnken, M. Schnölzer, A. Ciolfi, C. Di Stefano, M. Biffoni, L. Ricci-Vitiani, R. Pallini, R. De Maria, Integrin α7 Is a Functional Marker and Potential Therapeutic Target in Glioblastoma, Stem Cell 21, 35–50.e9 (2017).

9. S. Bao, Q. Wu, R. E. Mclendon, Y. Hao, Q. Shi, A. B. Hjelmeland, M. W. Dewhirst, D. D. Bigner, J. N. Rich, Glioma stem cells promote radioresistance by preferential activation of the DNA damage response, Nature 444, 756–760 (2006).

10. J. D. Lathia, S. C. Mack, E. E. Mulkearns-Hubert, C. L. L. Valentim, J. N. Rich, Cancer stem cells in glioblastoma, Genes & Development 29, 1203–1217 (2015).

11. J. S. Hale, B. Otvos, M. Sinyuk, A. G. Alvarado, M. Hitomi, K. Stoltz, Q. Wu, W. Flavahan, B. Levison, M. L. Johansen, D. Schmitt, J. M. Neltner, P. Huang, B. Ren, A. E. Sloan, R. L. Silverstein, C. L. Gladson, J. A. DiDonato, J. M. Brown, T. McIntyre, S. L. Hazen, C. Horbinski, J. N. Rich, J. D. Lathia, Cancer Stem Cell-Specific Scavenger Receptor CD36 Drives Glioblastoma Progression, Stem Cells 32, 1746–1758 (2014).

12. X. Wang, K. Yang, Q. Xie, Q. Wu, S. C. Mack, Y. Shi, L. J. Y. Kim, B. C. Prager, W. A. Flavahan, X. Liu, M. Singer, C. G. Hubert, T. E. Miller, W. Zhou, Z. Huang, X. Fang, A. Regev, M. L. S. agrave, T. H. Hwang, J. W. Locasale, S. Bao, J. N. Rich, Purine synthesis promotes maintenance of brain tumor initiating cells in glioma, Nat Neurosci, 1–15 (2017).

13. J. Jung, L. J. Y. Kim, X. Wang, Q. Wu, T. Sanvoranart, C. G. Hubert, B.C. Prager, L. C. Wallace, X. Jin, S. C. Mack, J. N. Rich, Nicotinamide metabolism regulates glioblastoma stem cell maintenance, JCI Insight 2, 766–24 (2017).

14. L. B. Hoang Minh, F. A. Siebzehnrubl, C. Yang, S. Suzuki Hatano, K. Dajac, T. Loche, N. Andrews, M. Schmoll Massari, J. Patel, K. Amin, A. Vuong, A. Jimenez Pascual, P. Kubilis, T. J. Garrett, C. Moneypenny, C. A. Pacak, J. Huang, E. J. Sayour, D. A. Mitchell, M. R. Sarkisian, B. A. Reynolds, L.P. Deleyrolle, Infiltrative and drug-resistant slow-cycling cells support metabolic heterogeneity in glioblastoma, The EMBO Journal, e98772–21 (2018).

15. R. Chianese, R. Coccurello, A. Viggiano, M. Scafuro, M. Fiore, G. Coppola, F. F. Operto, S. Fasano, S. Laye, R. Pierantoni, R. Meccariello, Impact of Dietary Fats on Brain Functions, Curr Neuropharmacol 16, 1059–1085 (2018).

16. C. He, D. Cheng, C. Peng, Y. Li, Y. Zhu, N. Lu, High-Fat Diet Induces Dysbiosis of Gastric Microbiota Prior to Gut Microbiota in Association With Metabolic Disorders in Mice, Front. Microbiol. 9, 979–9 (2018).

17. R. Mehrian-Shai, J. K. V. Reichardt, C. C. Harris, A. Toren, The Gut–Brain Axis, Paving the Way to Brain Cancer, TRENDS in CANCER, 1–8 (2019).

18. R. W. Crevel, J. V. Friend, B. F. Goodwin, W. E. Parish, High-fat diets and the immune response of C57Bl mice, Br. J. Nutr. 67, 17–26 (1992).

19. R. Wang, Physiological Implications of Hydrogen Sulfide: A Whiff Exploration That Blossomed, Physiol. Rev. 92, 791–896 (2012).

20. M. T. Peh, A. B. Anwar, D. S. W. Ng, M. S. Bin Mohamed Atan, S. D. Kumar, P. K. Moore, Effect of feeding a high fat diet on hydrogen sulfide (H2S) metabolism in the mouse, Nitric Oxide 41, 138–145 (2014).

21. B. D. Paul, S. H. Snyder, H2S: A Novel Gasotransmitter that Signals by Sulfhydration, Trends in Biochemical Sciences 40, 687–700 (2015).

22. D. Zhang, J. Du, C. Tang, Y. Huang, H. Jin, H2S-Induced Sulfhydration: Biological Function and Detection Methodology, Front. Pharmacol. 8, 814–13 (2017).

23. C. Szabo, C. Coletta, C. Chao, K. Módis, B. Szczesny, A. Papapetropoulos, M. R. Hellmich, Tumor-derived hydrogen sulfide, produced by cystathionine-β-synthase, stimulates bioenergetics, cell proliferation, and angiogenesis in colon cancer, Proceedings of the National Academy of Sciences 110, 12474–12479 (2013).

24. M. R. Hellmich, C. Szabo, Hydrogen Sulfide and Cancer, Handb Exp Pharmacol 230, 233–241 (2015).

25. D. Wu, W. Si, M. Wang, S. Lv, A. Ji, Y. Li, Hydrogen sulfide in cancer: Friend or foe? Nitric Oxide 50, 38–45 (2015).

26. N. Takano, Y. Sarfraz, D. M. Gilkes, P. Chaturvedi, L. Xiang, M. Suematsu, D. Zagzag, G. L. Semenza, Decreased Expression of Cystathionine-Synthase Promotes Glioma Tumorigenesis, Molecular Cancer Research 12, 1398–1406 (2014).

27. M. Wróbel, J. Czubak, P. Bronowicka-Adamska, H. Jurkowska, D. Adamek, B. Papla, Is Development of High-Grade Gliomas Sulfur-Dependent? Molecules 19, 21350–21362 (2014).

28. G. R. Villa, J. J. Hulce, C. Zanca, J. Bi, S. Ikegami, G. L. Cahill, Y. Gu, K. M. Lum, K. Masui, H. Yang, X. Rong, C. Hong, K. M. Turner, F. Liu, G. C. Hon, D. Jenkins, M. Martini,A. M. Armando, O. Quehenberger, T. F. Cloughesy, F. B. Furnari, W. K. Cavenee, P. Tontonoz, T. C. Gahman, A. K. Shiau, B. F. Cravatt, P. S. Mischel, An LXR-Cholesterol Axis Creates a Metabolic Co- Dependency for Brain Cancers, Cancer Cell 30, 683–693 (2016).

29. R. T. Martuscello, V. Vedam-Mai, D. J. McCarthy, M. E. Schmoll, M. A. Jundi, C. D. Louviere, B. G. Griffith, C. L. Skinner, O. Suslov, L. P. Deleyrolle, B. A. Reynolds, A Supplemented High-Fat Low-Carbohydrate Diet for the Treatment of Glioblastoma, Clin Cancer Res 22, 2482–2495 (2016).

30. J. Bi, S. Chowdhry, S. Wu, W. Zhang, K. Masui, P. S. Mischel, Altered cellular metabolism in gliomas — an emerging landscape of actionable co-dependency targets, Nature Reviews Cancer, 1–14 (2019).

31. E. E. Calle, C. Rodriguez, K. Walker-Thurmond, M. J. Thun, Overweight, obesity, and mortality from cancer in a prospectively studied cohort of U.S. adults, N Engl J Med 348, 1625–1638 (2003).

32. B. Lauby-Secretan, C. Scoccianti, D. Loomis, Y. Grosse, F. Bianchini, K. Straif, International Agency for Research on Cancer Handbook Working Group, Body Fatness and Cancer--Viewpoint of the IARC Working Group, N Engl J Med 375, 794–798 (2016).

33. M. N. Roberts, M. A. Wallace, A. A. Tomilov, Z. Zhou, G. R. Marcotte, D. Tran, G. Perez, E. Gutierrez-Casado, S. Koike, T. A. Knotts, D. M. Imai, S. M. Griffey, K. Kim, K. Hagopian, M. Z. McMackin, F. G. Haj, K. Baar, G. A. Cortopassi, J. J. Ramsey, J. A. Lopez-Dominguez, A Ketogenic Diet Extends Longevity and Healthspan in Adult Mice, Cell Metabolism 26, 539–546.e5 (2017).

34. J. C. Newman, A. J. Covarrubias, M. Zhao, X. Yu, P. Gut, C.-P. Ng, Y. Huang, S. Haldar, E. Verdin, Ketogenic Diet Reduces Midlife Mortality and Improves Memory in Aging Mice, Cell Metabolism 26, 547–557.e8 (2017).

35. L. M. Obeid, C. M. Linardic, L. A. Karolak, Y. A. Hannun, Programmed cell death induced by ceramide, Science 259, 1769–1771 (1993).

36. T. Yabu, H. Shiba, Y. Shibasaki, T. Nakanishi, S. Imamura, K. Touhata, M. Yamashita, Stress-induced ceramide generation and apoptosis via the phosphorylation and activation of nSMase1 by JNK signaling, Cell Death Differ. 22, 258–273 (2015).

37. A. G. Alvarado, P. S. Thiagarajan, E. E. Mulkearns-Hubert, D. J. Silver, J. S. Hale, T. Alban, S. M. Turaga, A. Jarrar, O. Reizes, M. S. Longworth, M. A. Vogelbaum, J. D. Lathia, Glioblastoma Cancer Stem Cells Evade Innate Immune Suppression of Self-Renewal through Reduced TLR4 Expression, Stem Cell, 1–17 (2016).

38. D. J. Silver, J. D. Lathia, Revealing the glioma cancer stem cell interactome, one niche at a time, J. Pathol. 244, 260–264 (2018).

39. E. A. Stoll, R. Makin, I. R. Sweet, A. J. Trevelyan, S. Miwa, P. J. Horner, D. M. Turnbull, Neural Stem Cells in the Adult Subventricular Zone Oxidize Fatty Acids to Produce Energy and Support Neurogenic Activity, Stem Cells 33, 2306–2319 (2015).

40. S. Yoshimoto, T. M. Loo, K. Atarashi, H. Kanda, S. Sato, S. Oyadomari, Y. Iwakura, H. Oshima, H. Morita, M. Hattori, K. Honda, Y. Ishikawa, E. Hara, N. Ohtani, Obesity-induced gut microbial metabolite promotes liver cancer through senescence secretome, Nature, 1–8 (2013).

41. T. N. Sergentanis, G. Tsivgoulis, C. Perlepe, I. Ntanasis-Stathopoulos, I.-G. Tzanninis, I. N. Sergentanis, T. Psaltopoulou, A. B. Hjelmeland, Ed. Obesity and Risk for Brain/CNS Tumors, Gliomas and Meningiomas: A Meta-Analysis, PLoS ONE 10, e0136974–29 (2015).

42. L. Disney-Hogg, A. Sud, P. J. Law, A. J. Cornish, Ben Kinnersley, Q. T. Ostrom, K. Labreche, J. E. Eckel-Passow, G. N. Armstrong, E. B. Claus, D. Il’yasova, J. Schildkraut, J. S. Barnholtz-Sloan, S. H. Olson, J. L. Bernstein, R. K. Lai, A. J. Swerdlow, M. Simon, P. Hoffmann, M. M. Nöthen, K.-H. Jöckel, S. Chanock, P. Rajaraman, C. Johansen, R. B. Jenkins, B. S. Melin, M. R. Wrensch, M. Sanson, M. L. Bondy, R. S. Houlston, Influence of obesity-related risk factors in the aetiology of glioma, Br J Cancer, 1–8 (2018).

43. A. S. Kuan, J. Green, C. M. Kitahara, A. B. De González, T. Key, G. K Reeves, S. Floud, A. Balkwill, K. Bradbury, L. M. Liao, N. D. Freedman, V. Beral, S. Sweetland, The Million Women Study, the NIH-AARP study, and the PLCO study, Diet and risk of glioma: combined analysis of 3 large prospective studies in the UK and USA, Neuro-Oncology 21, 944–952 (2019).

44. D. Hambardzumyan, Y.-K. Cheng, H. Haeno, E. C. Holland, F. Michor, Z. Su, Ed. The Probable Cell of Origin of NF1- and PDGF-Driven Glioblastomas, PLoS ONE 6, e24454 (2011).

45. J. H. Lee, J. E. Lee, J. Y. Kahng, S. H. Kim, J. S. Park, S. J. Yoon, J.-Y. Um, W. K. Kim, J.-K. Lee, J. Park, E. H. Kim, J.-H. Lee, J.-H. Lee, W.-S. Chung, Y. S. Ju, S.-H. Park, J. H. Chang, S.-G. Kang, J. H. Lee, Human glioblastoma arises from subventricular zone cells with low-level driver mutations, Nature, 1–24 (2018).

46. P. Mukherjee, Z. M. Augur, M. Li, C. Hill, B. Greenwood, M. A. Domin, G. Kondakci, N. R. Narain, M. A. Kiebish, R. T. Bronson, G. Arismendi-Morillo, C. Chinopoulos, T. N. Seyfried, Therapeutic benefit of combining calorie-restricted ketogenic diet and glutamine targeting in late-stage experimental glioblastoma, Communications Biology, 1–14 (2019).

47. X. Wang, K. Yang, Q. Wu, L. J. Y. Kim, A. R. Morton, R. C. Gimple, B. C. Prager, Y. Shi, W. Zhou, S. Bhargava, Z. Zhu, L. Jiang, W. Tao, Z. Qiu, L. Zhao, G. Zhang, X. Li, S. Agnihotri, P. S. Mischel, S. C. Mack, S. Bao, J. N. Rich, Targeting pyrimidine synthesis accentuates molecular therapy response in glioblastoma stem cells, Science Translational Medicine 11, eaau4972 (2019).

48. C. Hine, E. Harputlugil, Y. Zhang, C. Ruckenstuhl, B. C. Lee, L. Brace, A. Longchamp, J. H. Treviño-Villarreal, P. Mejia, C. K. Ozaki, R. Wang, V. N. Gladyshev, F. Madeo, W. B. Mair, J. R. Mitchell, Endogenous Hydrogen Sulfide Production Is Essential for Dietary Restriction Benefits, Cell 160, 132–144 (2015).

49. C. Hine, Y. Zhu, A. N. Hollenberg, J. R. Mitchell, Dietary and Endocrine Regulation of Endogenous Hydrogen Sulfide Production: Implications for Longevity, Antioxidants & Redox Signaling 28, 1483–1502 (2018).

50. A. D. Gromovsky, R. C. Schugar, A. L. Brown, R. N. Helsley, A. C. Burrows, D. Ferguson, R. Zhang, B. E. Sansbury, R. G. Lee, R. E. Morton, D. S. Allende, J. S. Parks, M. Spite, J. M. Brown, ∆-5 Fatty Acid Desaturase FADS1 Impacts Metabolic Disease by Balancing Proinflammatory and Proresolving Lipid Mediators, Arterioscler. Thromb. Vasc. Biol. 38, 218–231 (2018).

51. C. K. A. Neumann, D. J. Silver, V. Venkateshwari, R. Zhang, C. A. Traughber, C. Przybycin, D. Bayik, J. D. Smith, J. D. Lathia, B. I. Rini, J. M. Brown, MBOAT7-driven phosphatidylinositol remodeling promotes the progression of clear cell renal carcinoma, Molecular Metabolism 34, 136–145 (2020).

52. Y. Hu, G. K. Smyth, ELDA: Extreme limiting dilution analysis for comparing depleted and enriched populations in stem cell and other assays, Journal of Immunological Methods 347, 70–78 (2009).

53. N. Bithi, C. Link, R. Wang, B. Willard, C. Hine. BioRχiv, 2019, Dietary restriction transforms the protein sulfhydrome in a tissue-specific and cystathionine γ-lyase-dependent manner, biorxiv.org, doi:10.1101/869271.

54. M. Goldman, B. Craft, A. Brooks, J. Zhu, D. Haussler. BioRχiv, 2018, The UCSC Xena Platform for cancer genomics data visualization and interpretation, biorxiv.org, doi:10.1101/326470.

